# MMEJ-based Precision Gene Editing for applications in Gene Therapy and Functional Genomics

**DOI:** 10.1101/2020.04.25.060541

**Authors:** Gabriel Martínez-Gálvez, Armando Manduca, Stephen C. Ekker

## Abstract

Experiments in gene editing commonly elicit error-prone non-homologous end joining for DNA double-strand break (DSB) repair. Microhomology-mediated end joining (MMEJ) can generate more predictable outcomes for functional genomic and somatic therapeutic applications. MENTHU is a computational tool that predicts nuclease-targetable sites likely to result in MMEJ-repaired, homogeneous genotypes (PreMAs) in zebrafish. We deployed MENTHU on 5,885 distinct Cas9-mediated DSBs in mouse embryonic stem cells, and compared the predictions to those by inDelphi, another DSB repair predictive algorithm. MENTHU correctly identified 46% of all PreMAs available, doubling the sensitivity of inDelphi. We also introduce MENTHU@4, an MENTHU update trained on this large dataset. We trained two MENTHU-based algorithms on this larger dataset and validated them against each other, MENTHU, and inDelphi. Finally, we estimated the frequency and distribution of SpCas9-targetable PreMAs in vertebrate coding regions to evaluate MMEJ-based targeting for gene discovery. 44 out of 54 genes (81%) contained at least one early out-of-frame PreMA and 48 out of 54 (89%) did so when also considering Cas12a. We suggest that MMEJ can be deployed at scale for reverse genetics screenings and with sufficient intra-gene density rates to be viable for nearly all loss-of-function based gene editing therapeutic applications.

## INTRODUCTION

Precision in gene editing is currently limited by the high variability in genotypic outcomes of the commonly deployed NHEJ repair pathway or the low efficiency of the more precise homologous recombination pathway (for reviews see (1-4)). These shortcomings often result in complicated and labor-intensive selection processes for identifying gene edits of interest, particularly if pursuing bi-allelic editing of vertebrate cells (5,6). These limitations of NHEJ-based gene editing also potentially reduce its utility in somatic applications such as gene therapy or gene discovery. To address this technical gap in the field, we have developed alternative gene editing approaches that aim to elicit MMEJ (Microhomology-Mediated End Joining) instead of NHEJ and are both precise and efficient and suitable for reverse genetics applications.

As opposed to NHEJ, MMEJ is thought to bridge a DSB (DNA double-strand break) by annealing a pair of short stretches (3-12 bases) of single-stranded cis-homologies (microhomologies: µHs) exposed by the 5’-resection of the DSB ends (7), the trimming of the resulting 3’-flap overhangs (8,9), and finally a DNA ligation (10,11). This process results in a characteristic deletion where the sequence between the pair of µHs used for repair and one of the repeats itself is lost (12) (see (13) for a review). This deletion pattern is then useful as a heuristic to identify probable MMEJ-based repairs from a mixed-repair pool. The ability to generate predictable genotypes (14), sometimes even resulting in an identical allele in over half of the editing outcomes (15), makes MMEJ an attractive alternative to NHEJ for precision genome engineering (16-19).

We and others (15,16,19) have shown that DSBs directed at sites likely to be repaired via MMEJ significantly increase the homogeneity of the resulting repair outcomes in zebrafish. Despite there being no known clear biochemical mechanism as to how MMEJ repairs DSBs, we and others (20-23) have published software tools that predict the occurrence of MMEJ based on the sequence context surrounding any given target DSB site. In particular, we have published MENTHU (22), which screens genetic sequences for DSBs likely to generate consistent repair outcomes (with 50% or more of a DSB repair profile displaying the same genotype) as a result of MMEJ, aka PreMAs (predominant MMEJ alleles) (Figure 1A and 1B). As opposed to conventional targeting designs, using MENTHU-recommended SpCas9 and TALEN cut sites resulted in PreMAs more often, facilitating subsequent zebrafish mutant screenings by decreasing the sequence variability of the resulting allele pool (15). In parallel, the tools ForeCasT (20) and inDelphi (23) were simultaneously and independently developed to predict the probability of occurrence of individual repair outcomes after Cas9-mediated DSBs on mammalian cells, and were shortly followed by Lindel (21). Here, we provide an experimental assessment of MENTHU in an independent gene editing dataset to evaluate the predictive performance of MENTHU on mammalian cells, compare it with that of inDelphi, suggest improvements to MENTHU, and assess the practicality of PreMA predictions prospectively in a use-case scenario.

**Figure 1.**
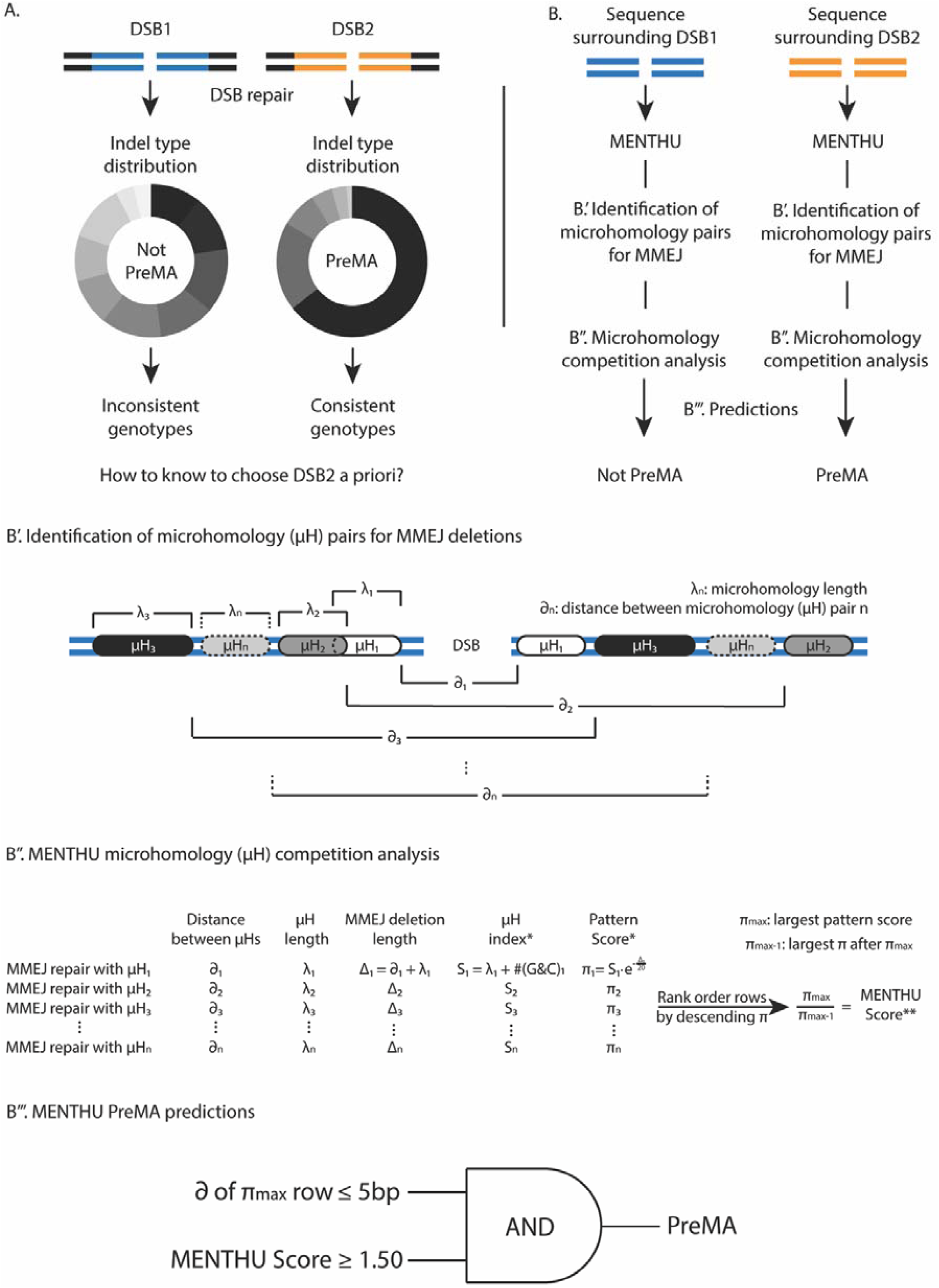
MENTHU elucidates which DNA double-strand breaks likely result in more consistent genotypes. A. Different DNA double-strand breaks (DSBs) can generate indel profiles with dissimilar distributions. Being able to discern the genotype heterogeneity level between targetable DSBs prior to experimental applications would be beneficial for reverse genetics and gene therapy applications. B. MENTHU (22) is a software tool that analyzes the DNA sequence surrounding any given DSB and predicts whether it will result in a PreMA: an MMEJ-mediated repaired sequence where half or more of the repair outcomes share the same genotype. B’. MENTHU identifies every possible µH pair (with homology arms µH_1_ to µH_n_ of length λ_1_ to λ_n_) and calculates the corresponding distance between the µHs of each pair (*∂*_1_ to *∂*_n_). B’’. Based on the expected MMEJ deletion pattern, *∂*_i_ and λ_i_ are used to calculate the expected deletion length Δ_i_. Pattern scores π_i_ for every possible MMEJ deletion are calculated as described by Bae et al. (16). The MMEJ deletions are then rank ordered by descending pattern score and a MENTHU Score for the DSB is calculated by taking the ratio between the largest π_max_ and the second largest pattern score π_max-1_. B’’’ MENTHU utilizes two criteria that need to be concomitantly true for a DSB to be labeled as a PreMA. The *∂* of the MMEJ-deletion with the highest pattern score πmax and the MENTHU Score for the DSB need to be less than or equal to 5bp and more than or equal to 1.50, respectively, for a positive PreMA prediction.

## MATERIAL AND METHODS

### Sequence data acquisition, inclusion, and classification

Due to the lack of a comprehensive database for deep sequencing results of DSB repair events, a subset of the data generated by Allen and collaborators (20) was chosen to assess the predictive performance of MENTHU and compare it to inDelphi predictions. A total of 41,388 different Cas9-mediated gene edits of mESC cells were downloaded from https://figshare.com/articles/processed_mutational_profiles/7312067. For each DSB event, all resulting repair sequences from all experimental replicates were combined into a single pool, consolidated by sequence, and rank ordered by number of reads. Subsequently, the most frequently observed read was aligned to its corresponding WT sequence (obtained from the Supplementary Data 1 at (20)) using the pairwiseAlignment function from the Bioconductor Biostrings package (version 2.54.0) with a substitution matrix that penalized mismatches heavily (match = 1, mismatch = -50). Only those alignments that could be explained by a single, simple indel (insertion or deletion) were included in the analysis, since MENTHU and inDelphi were trained to predict, respectively, single-deletion and single-indel repairs exclusively. Each of these were classified into one of four different groups based on the nature of the observed indel: a 1bp or >1bp insertion, and an MMEJ or non-MMEJ deletion. Following the proposed mechanism for MMEJ repair, MMEJ deletions were defined as those that displayed, in the WT sequence, two µHs of at least 3bp at each side of the expected DSB, but later collapsed into one in the repaired read, deleting any intervening nucleotides. Deletions that did not follow this pattern were considered non-MMEJ deletions. Lastly, since the original Allen et al. data (20) included many gRNAs targeting artificially manufactured targets, only those that targeted genomic sites were included in the analysis so the results of this study would be representative of genome-targeting experiments. This process culminated in a total of 5,885 Cas9-mediated edits, each with its WT sequence, most frequent repair sequence, and its corresponding observed frequency (Figure 2A).

**Figure 2.**
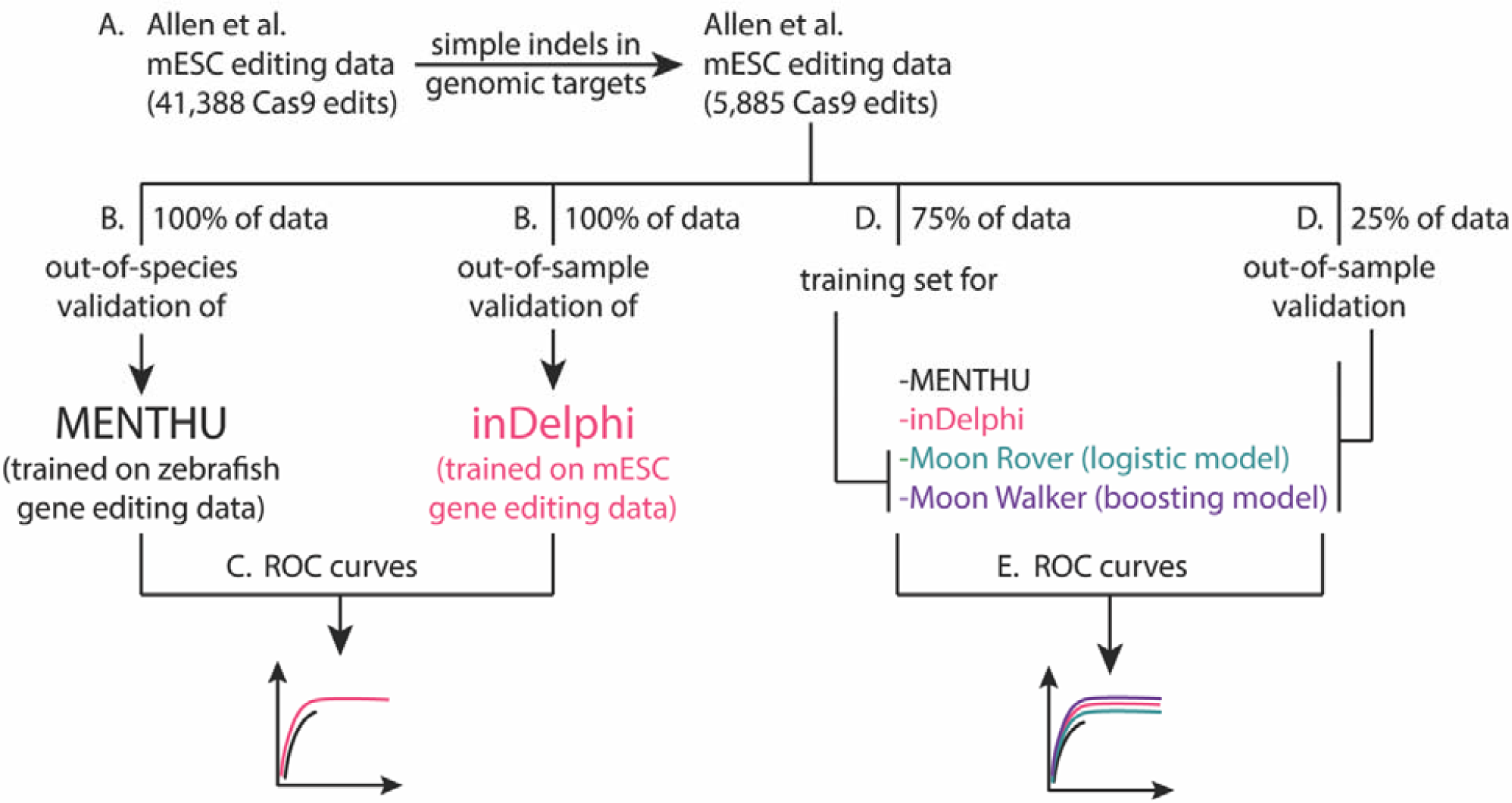
Workflow of the independent assessment of the ability of MENTHU to predict PreMAs. A. A large gene editing dataset was filtered to only include genomic DSB repair outcomes that resulted in simple indels (i.e. resulting in single deletions or insertions). B. This dataset was used to assess the viability of MENTHU PreMA predictions in a mammalian cell system (mouse ESC cells [mESCs]), since MENTHU was originally validated in zebrafish embryos. To contextualize any MENTHU claims, the same dataset was used to generate PreMA predictions using inDelphi, a similar-purpose software tool in the recent literature validated in mESC cells. C. Receiving operating characteristic (ROC) curves were used to compare the ability to predict PreMAs by MENTHU and inDelphi. D. To investigate whether the MENTHU prediction scheme maximizes the predictive capacity of the features it uses for classification, the large dataset described in (A) was split into 75% for the training of machine learning models for PreMA predictions and 25% for the out-of-sample evaluation of these models. E. The training set in (D) was used to train Moon Rover (a logistic regression classifier) and Moon Walker (a gradient boosting machine classifier). ROC curves for Moon Rover and Moon Walker were generated based on their predictive performance on the testing set in (D), and were plotted together with ROC curves of MENTHU and inDelphi on the same testing set for reference.

### Comparison between PreMA predictive performance of MENTHU and inDelphi

For each for the 5,885 edits, a 52bp sequence window centered at the Cas9 expected DSB location (i.e. 3 bases upstream of NGG PAM) was extracted. These short sequences served as inputs for the command-line versions of MENTHU (R) and inDelphi (python 2.7) to generate PreMA predictions. For every input, MENTHU outputs a data frame with all possible MMEJ-based deletions within the 52bp sequence (Figure 1B’) and rank orders them by MENTHU score, as described in Mann et al. (22). We classified MENTHU predictions as recommended by Ata et al. (15), labelling as PreMA any site displaying (a) a MENTHU score of 1.50 or above and (b) a distance of 5bp or less in the WT sequence between the µHs utilized for MMEJ of the most frequent predicted repair outcome (Figure 1B’’’). On the other hand, inDelphi outputs the probability of occurrence of a list of potential repair outcomes for every input. Consequently, any site where the most likely prediction has a probability of 0.50 or more and displays the MMEJ deletion pattern was classified as a PreMA. PreMA predictions from both MENTHU and inDelphi were compared to their corresponding ground truth sequencing data (Figure 2B) and classified as true positives (TP, when a PreMA prediction matched the data), true negatives (TN, when a non-PreMA prediction matched the data), false positives (FP, when a PreMA prediction did not match the data), and false negatives (FN, when a non-PreMA prediction did not match the data). Sensitivity (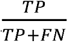 : the percentage of actual PreMAs correctly classified as such), specificity (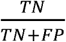 : the percentage of actual non-PreMAs correctly classified as such), and positive predictive value (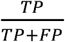 : PPV, percentage of correct positive PreMA predictions) were calculated for both MENTHU and inDelphi. The possibility of utilizing these tools synergistically was assessed by calculating sensitivity, specificity, and PPV after taking the initial PreMA predictions for both tools and labeling a site as a PreMA either when MENTHU and inDelphi agreed, or when either one of them predicted a PreMA individually.

### Generation of receiving operating characteristic curves for MENTHU and inDelphi

Receiving operating characteristic (ROC) curves are a standard technique for evaluating binary classifiers (24), and are plots of TP (sensitivity) vs. FP (1-specificity) as a function of varying classification thresholds. ROC curves were generated for MENTHU and inDelphi (Figure 2C) to evaluate their predictive performance independent of specific values for model thresholds (i.e. MENTHU score and inDelphi probability). The MENTHU ROC curve was generated by varying the MENTHU score classification threshold from 0 to infinity while leaving the µH distance requirement (less than or equal to 5bp) constant. ROC curves for inDelphi were generated by varying the predicted probability threshold used for PreMA classifications (originally 0.50).

### Development of MENTHU-based PreMA prediction models

The original MENTHU, as described by Ata et al. (15), is a threshold-based, two-feature PreMA prediction scheme (Figure 1B’’’). To investigate the impact of the distance threshold component and complement the ROC curve analysis described above, we calculated the sensitivity, specificity, and PPV values of PreMA predictions at 3 other threshold values (less than or equal to 3, 4, and 6bp: MENTHU@3, MENTHU@4, and MENTHU@6, respectively) while keeping the MENTHU score threshold constant. Additionally, we examined the impact of combining the prediction outcomes of the original MENTHU, MENTHU@4, and inDelphi.

The dataset from Allen et al. (20) was also used to train and test two machine learning models (Moon Rover and Moon Walker) utilizing the same features as MENTHU uses to predict PreMAs (i.e. the MENTHU score and the distance between the µH pair for the top predicted MMEJ-based outcome). Significant improvements of either of these models over the original MENTHU would suggest a better way to utilize these two features to improve predictive performance. The 5,885 data points from Allen et al. (20) were divided into a training set and a test set in a 70-30% split (4,120 and 1,765 respectively), with the PreMA to non-PreMA ratio remaining constant in both sets (Figure 2D). Moon Rover is a logistic regression model and Moon Walker is a gradient boosting model (25). The latter used a 10-fold cross validation scheme to choose the set of model hyperparameters that displayed the highest ROC curve area under the curve. Each hyperparameter set was defined by a grid-search of the number of boosting iterations (decision trees), maximum tree depth, minimum amount of observations per node, and shrinkage level (regularization constant). The model performance was assessed by making out-of-sample predictions on the test set. MENTHU and inDelphi ROC curves on the test set were also calculated for comparison (Figure 2D and 2E). Both models were trained and evaluated using the R-based caret package (26).

### Assessment of PreMA targeting for protein knock-outs

Accurate prediction of PreMAs would be of limited value if their frequency of general localization within a gene were not useful experimentally. Therefore, we investigated the intragenic density and localization of 28 human and 26 zebrafish genes (Table 1) for PreMAs likely to induce protein knockout experiments. The first 30% of the cDNA sequence of each gene was screened for potential Cas9 and Cas12a-mediated PreMAs resulting in out-of-frame mutations using MENTHU. To ensure screening of at least 150bp per cDNA, the first 182bp were screened for shorter coding sequences (150bp + 32bp upstream of DSB for sequence context).

**Table 1.**
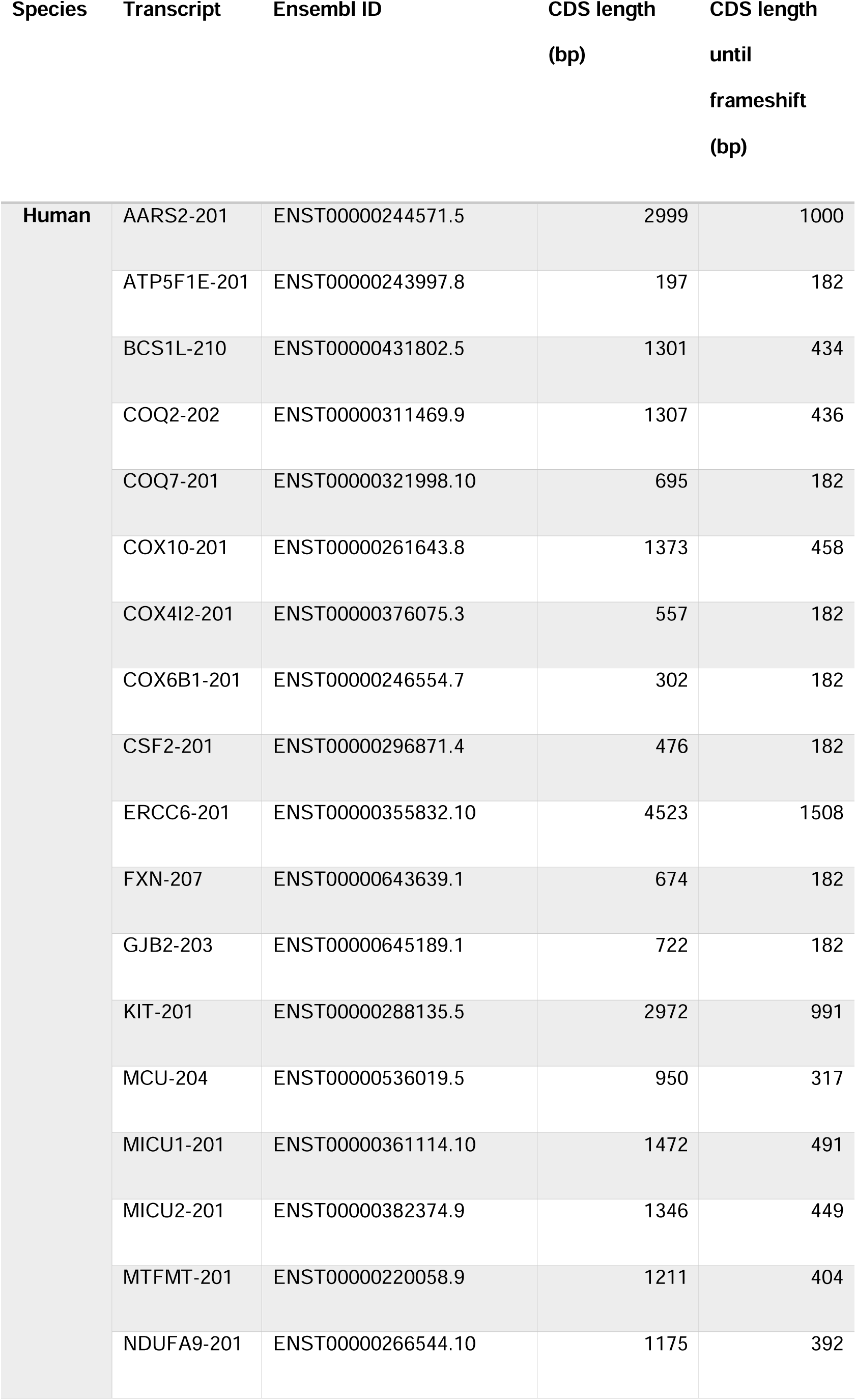
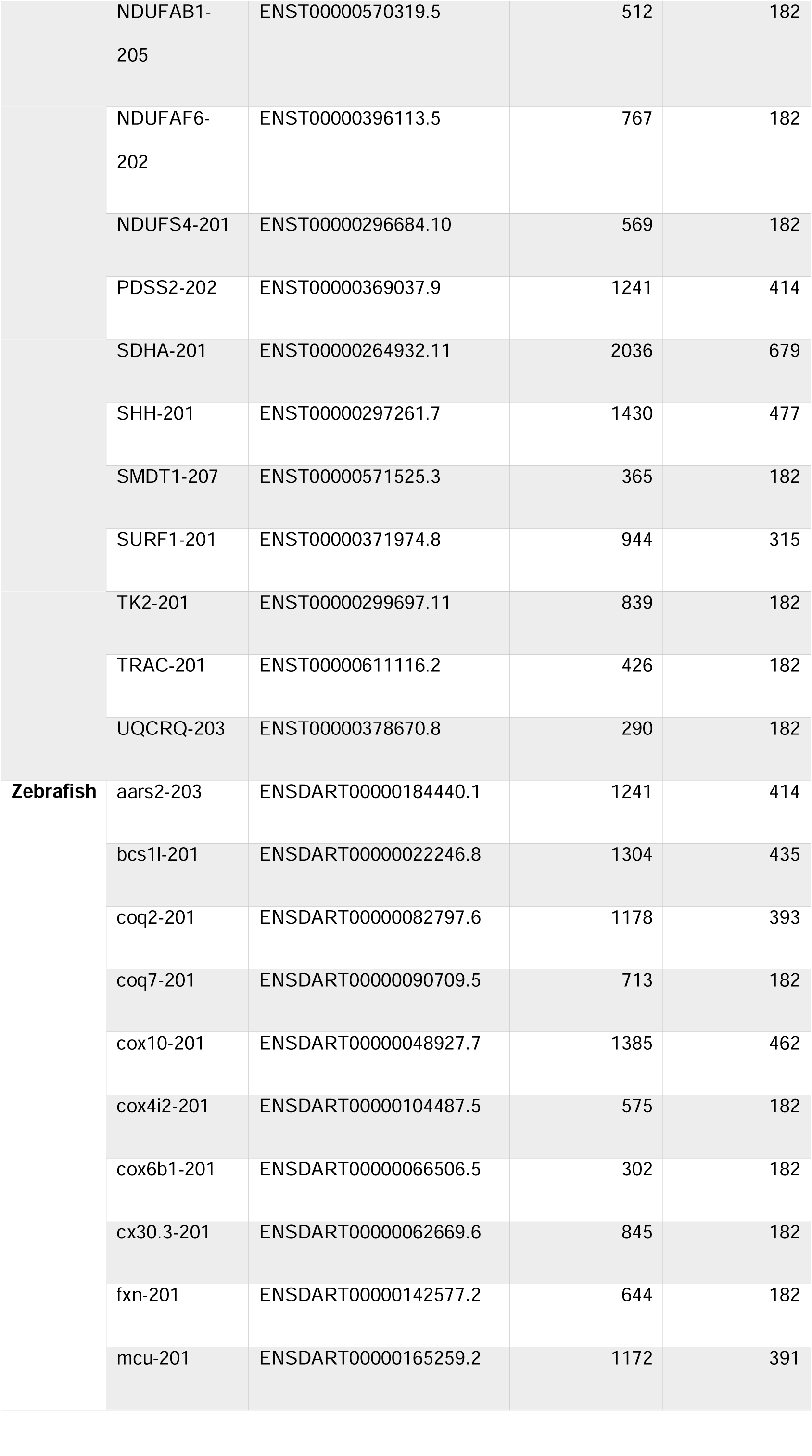
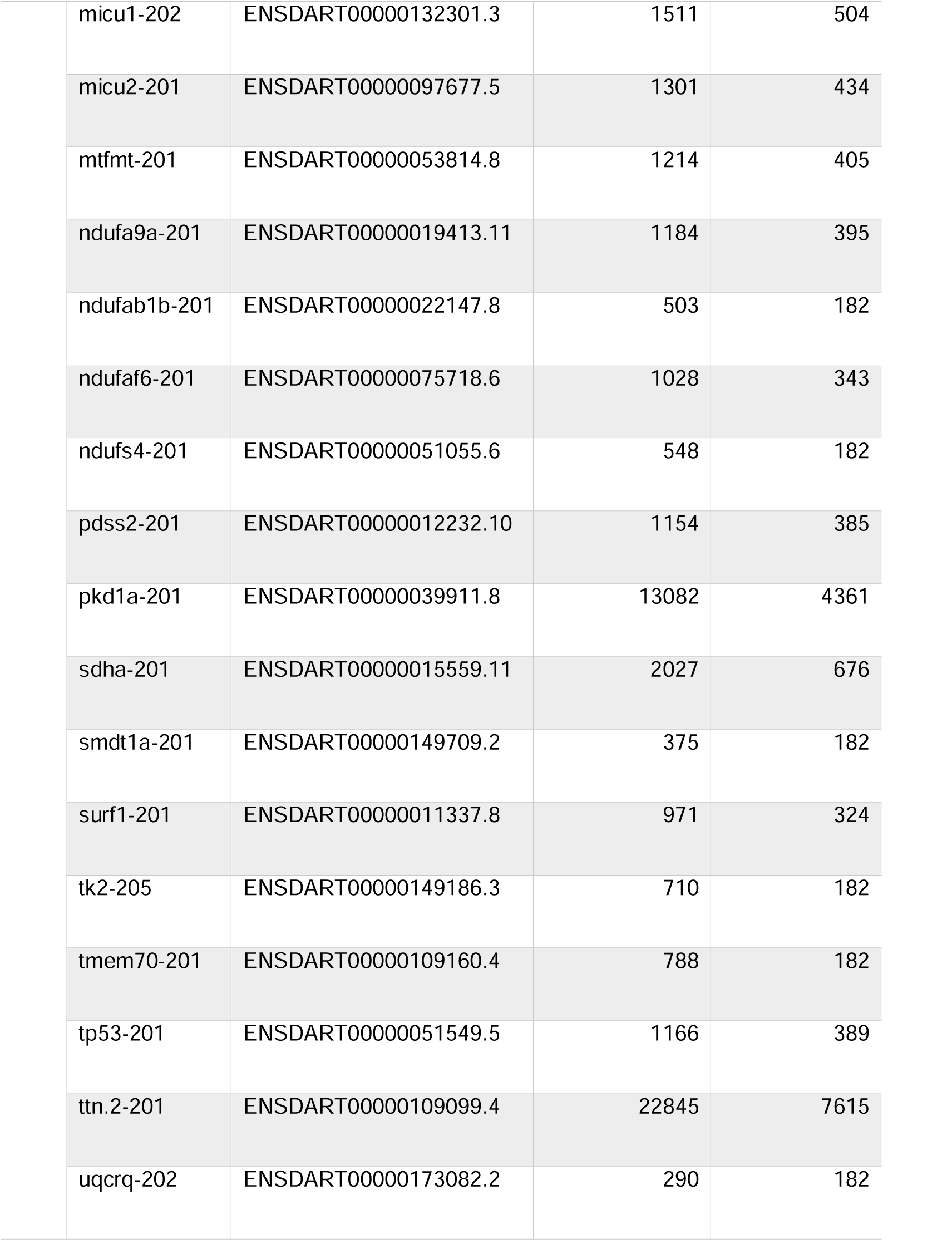
List of coding sequences screened for early frameshift PreMA alleles using MENTHU.

TABLE AND FIGURES LEGENDS
| Species | Transcript | Ensembl ID | CDS length<br>(bp) | CDS length<br>until<br>frameshift<br>(bp) |
| --- | --- | --- | --- | --- |
| Human | AARS2-201 | ENST00000244571.5 | 2999 | 1000 |
|  | ATP5F1E-201 | ENST00000243997.8 | 197 | 182 |
|  | BCS1L-210 | ENST00000431802.5 | 1301 | 434 |
|  | COQ2-202 | ENST00000311469.9 | 1307 | 436 |
|  | COQ7-201 | ENST00000321998.10 | 695 | 182 |
|  | COX10-201 | ENST00000261643.8 | 1373 | 458 |
|  | COX4I2-201 | ENST00000376075.3 | 557 | 182 |
|  | COX6B1-201 | ENST00000246554.7 | 302 | 182 |
|  | CSF2-201 | ENST00000296871.4 | 476 | 182 |
|  | ERCC6-201 | ENST00000355832.10 | 4523 | 1508 |
|  | FXN-207 | ENST00000643639.1 | 674 | 182 |
|  | GJB2-203 | ENST00000645189.1 | 722 | 182 |
|  | KIT-201 | ENST00000288135.5 | 2972 | 991 |
|  | MCU-204 | ENST00000536019.5 | 950 | 317 |
|  | MICU1-201 | ENST00000361114.10 | 1472 | 491 |
|  | MICU2-201 | ENST00000382374.9 | 1346 | 449 |
|  | MTFMT-201 | ENST00000220058.9 | 1211 | 404 |
|  | NDUFA9-201 | ENST00000266544.10 | 1175 | 392 |
|  | NDUFAB1-205 | ENST00000570319.5 | 512 | 182 |
|  | NDUFAF6-202 | ENST00000396113.5 | 767 | 182 |
|  | NDUFS4-201 | ENST00000296684.10 | 569 | 182 |
|  | PDSS2-202 | ENST00000369037.9 | 1241 | 414 |
|  | SDHA-201 | ENST00000264932.11 | 2036 | 679 |
|  | SHH-201 | ENST00000297261.7 | 1430 | 477 |
|  | SMDT1-207 | ENST00000571525.3 | 365 | 182 |
|  | SURF1-201 | ENST00000371974.8 | 944 | 315 |
|  | TK2-201 | ENST00000299697.11 | 839 | 182 |
|  | TRAC-201 | ENST00000611116.2 | 426 | 182 |
|  | UQCRQ-203 | ENST00000378670.8 | 290 | 182 |
| **Zebrafish** | aars2-203 | ENSDART00000184440.1 | 1241 | 414 |
|  | bcs1l-201 | ENSDART00000022246.8 | 1304 | 435 |
|  | coq2-201 | ENSDART00000082797.6 | 1178 | 393 |
|  | coq7-201 | ENSDART00000090709.5 | 713 | 182 |
|  | cox10-201 | ENSDART00000048927.7 | 1385 | 462 |
|  | cox4i2-201 | ENSDART00000104487.5 | 575 | 182 |
|  | cox6b1-201 | ENSDART00000066506.5 | 302 | 182 |
|  | cx30.3-201 | ENSDART00000062669.6 | 845 | 182 |
|  | fxn-201 | ENSDART00000142577.2 | 644 | 182 |
|  | mcu-201 | ENSDART00000165259.2 | 1172 | 391 |

|  |  |  |  |
| --- | --- | --- | --- |
| micu1-202 | ENSDART00000132301.3 | 1511 | 504 |
| micu2-201 | ENSDART00000097677.5 | 1301 | 434 |
| mtfmt-201 | ENSDART00000053814.8 | 1214 | 405 |
| ndufa9a-201 | ENSDART00000019413.11 | 1184 | 395 |
| ndufab1b-201 | ENSDART00000022147.8 | 503 | 182 |
| ndufaf6-201 | ENSDART00000075718.6 | 1028 | 343 |
| ndufs4-201 | ENSDART00000051055.6 | 548 | 182 |
| pdss2-201 | ENSDART00000012232.10 | 1154 | 385 |
| pkd1a-201 | ENSDART00000039911.8 | 13082 | 4361 |
| sdha-201 | ENSDART00000015559.11 | 2027 | 676 |
| smdt1a-201 | ENSDART00000149709.2 | 375 | 182 |
| surf1-201 | ENSDART00000011337.8 | 971 | 324 |
| tk2-205 | ENSDART00000149186.3 | 710 | 182 |
| tmem70-201 | ENSDART00000109160.4 | 788 | 182 |
| tp53-201 | ENSDART00000051549.5 | 1166 | 389 |
| ttn.2-201 | ENSDART00000109099.4 | 22845 | 7615 |
| uqcrq-202 | ENSDART00000173082.2 | 290 | 182 |

## RESULTS

### Data inclusion

We assembled a test set of Cas9 target-repair pairs from the Allen et al. data (20) that targeted mESC genomic sites and assembled the sequence outcomes from simple indels in the analysis described here. For modelling work in human DNA contexts, we focused on the 5,885 loci subset whose corresponding gRNAs target the human genome and did not display more than one indel outcome (Figure 2A). Based on the sequence characteristics of every individual alignment, these were divided, approximately, into 54% non-MMEJ deletions, 31% MMEJ deletions, 14% 1bp insertions, and 0.2% 1+bp insertions.

### Comparison between PreMA predictive performance of MENTHU and inDelphi

Sensitivity, specificity, and positive predictive value (PPV) were calculated for the MENTHU and inDelphi PreMA predictions of each of the 5,885 Cas9-mediated breaks. The corresponding confusion matrices are shown in Figure 3A. Of the 614 PreMAs in the data, MENTHU correctly identified twice the total number as inDelphi (with sensitivities of 46 and 23%, respectively), although with a lower correct classification-rate of PreMA-positive events (55 to 76% PPV, respectively) and slightly lower specificity (96 and 99%, respectively). In terms of PreMA prediction coverage, 27.0% of all available PreMAs were uniquely predicted by MENTHU, 4.6% by inDelphi only, and 18.9% by both (166, 28, and 116 out of 614, respectively). In contrast, the majority (95.3%) of the 5,271 non-PreMAs in the data were correctly classified by both tools. Further analysis into these differences revealed that 10/28 of the PreMAs found by inDelphi but not MENTHU failed the MENTHU µH proximity criterion (see *∂* at Figure 1B and “Sequence data acquisition, inclusion, and classification” and “Sequence data acquisition, inclusion, and classification” in Materials and Methods), and the remaining 18/28 failed the MENTHU Score 1.50 cutoff (four of them by 0.01 or less).

**Figure 3.**
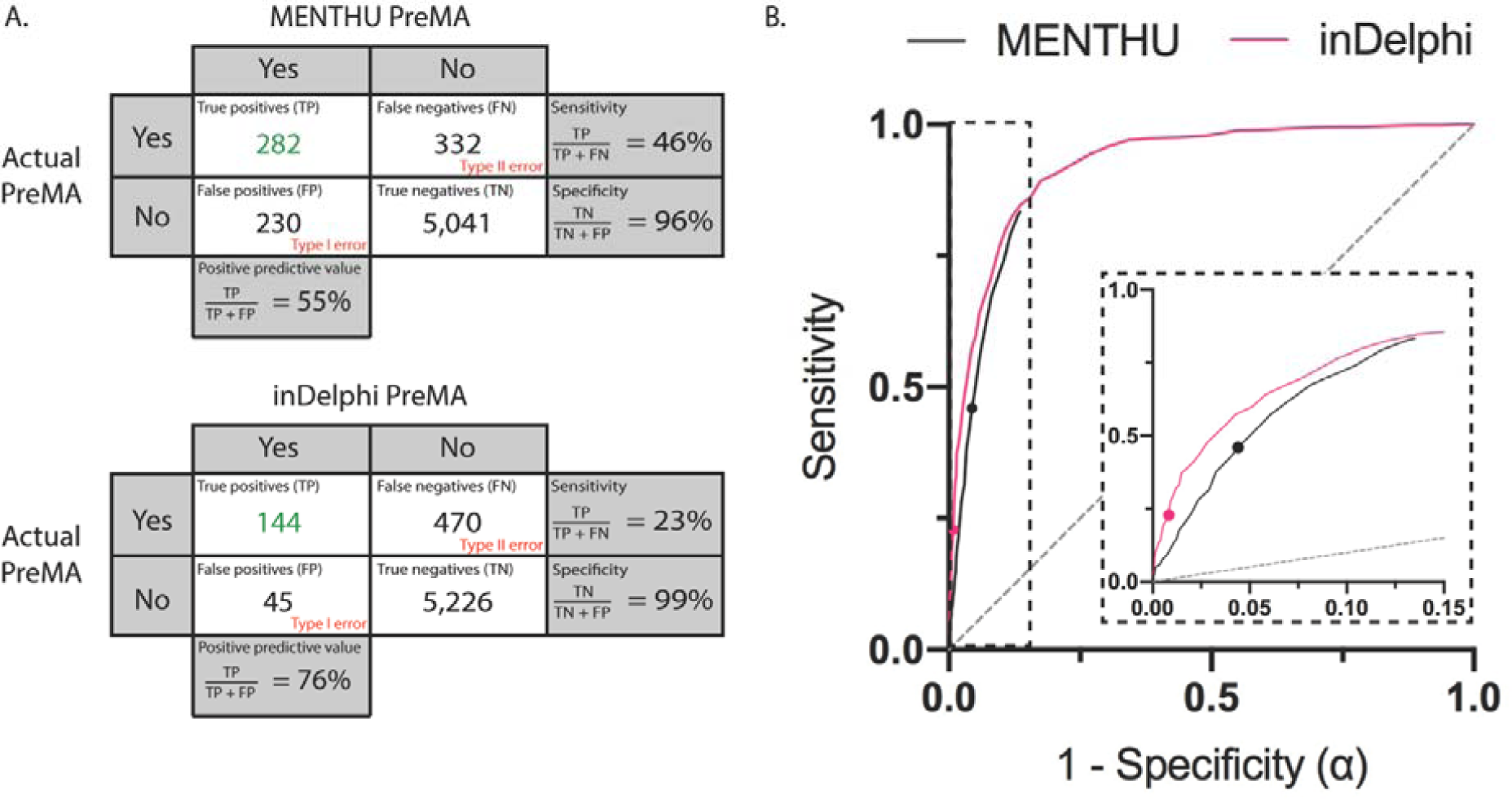
Comparison of the performance of the published versions of MENTHU and inDelphi in predicting PreMAs in a large, out-of-sample dataset. A. Confusion matrices for PreMA predictions by MENTHU (top) and inDelphi (bottom). Rows indicate the PreMA status of 5,885 Cas9 generated mutation profiles in mESC cells taken from Allen et al. (20). Columns denote the PreMA predictions by MENTHU and inDelphi and rows refer to the actual PreMA status. Sensitivity is the proportion of positive-PreMAs correctly predicted as such. Specificity is the proportion of negative-PreMAs correctly predicted as such. PPV is the proportion of correct predictions of positive-PreMAs. B. Receiving Operating Curves (ROCs) comparing MENTHU and inDelphi PreMA predictions. Here, sensitivity is plotted against 1 – specificity (or the probability of a type I error: α) as a function of varying prediction thresholds. The two points emphasized represent the published thresholds for both tools. The MENTHU ROC curve was generated by varying the MENTHU score threshold for PreMA classification. In the inDelphi ROC curve, the minimum threshold probability of the most frequent predicted read was varied. The MENTHU curve is truncated because its second classification criterium regarding the maximum distance between µHs allowed for MMEJ classification does not allow for a higher sensitivity. The inset is a blowup of the region where MENTHU is present.

A breakdown of the false-positive PreMA predictions by MENTHU revealed that over 60% of them failed because of not reaching the ≥ 50% frequency requirement, albeit displaying the characteristic MMEJ deletion pattern. Over half of these had greater than 40% frequency, and most of those greater than 46% frequency. Importantly, most of these MMEJ outcomes (97%) displayed exactly the sequence changes predicted by MENTHU. This latest finding was consistent across true positives by both MENTHU and inDelphi where, respectively, 100% and 99% of the sequence predictions matched the observed predominant sequence.

### Receiving operating characteristic (ROC) curves for MENTHU and inDelphi predictions

ROC curves for MENTHU and inDelphi are shown in Figure 3B. The MENTHU curve was plotted by varying the MENTHU score threshold utilized for PreMA classification. The MENTHU ROC curve is truncated since MENTHU classifies any MMEJ prediction that does not comply with the maximum of 5bp distance between µHs as a non-PreMA. Hence, its maximum achievable sensitivity was 83.39% (the top-left most point of the ROC curve).

### Development of MENTHU-based PreMA prediction models

As evidenced by the ROC curves described above, choosing different MENTHU score thresholds results in trade-offs between sensitivity and specificity. Since MENTHU classifications rely on two different thresholds (Figure 1B’’’), we looked at varying the distance threshold (*∂*) while keeping the MENTHU score threshold constant to observe changes in the predictive performance of PreMAs (Table 2). Figure 4A shows the change of MMEJ repair events as a function of *∂* and their corresponding PreMA distribution across all 1,844 MMEJ repaired events in the data. Figure 4B displays the distribution of PreMAs across each bin of Figure 4A. We found that, on this data set, MENTHU increased its PPV and specificity to approximately 65% and 97% (∼10% and a ∼1.5% increase, respectively) by decreasing the *∂* by 1bp (to 4bp) in exchange for a ∼4.5% sensitivity loss (MENTHU@4). We also looked at whether combining the MENTHUs and/or inDelphi PreMA predictions resulted in a better performance (Table 3). The best combination was MENTHU@4 or inDelphi (i.e., predict a PreMA if either algorithm makes this prediction), achieving a 64% PPV and a ∼1% increase in both sensitivity and specificity in comparison to the original MENTHU. In order to investigate if the way the two features that MENTHU uses for PreMA predictions can be optimized, two machine learning algorithms (Moon Rover and Moon Walker) were developed using the same inputs and outputs as MENTHU.

**Table 2.** Summary of the PreMA predictive performance of MENTHU at different µH distance thresholds in bp (@x bp). * MENTHU@5 representes the performance metrics by the original MENTHU. PPV: Positive predictive value, the percentage of correct positive PreMA predictions.

**Model Sensitivity Specificity PPV**
|  |  |  |  |
| --- | --- | --- | --- |
| *MENTHU@3* | 36.64% | 97.99% | 67.98% |
| *MENTHU@4* | 41.53% | 97.36% | 64.72% |
| *\*MENTHU@5* | <u>45.93%</u> | <u>95.64%</u> | <u>55.08%</u> |
| *MENTHU@6* | 48.70% | 93.54% | 47.16% |

**Table 3.** Summary of the PreMA predictive performance of different combinations between MENTHU and inDelphi. PPV: Positive predictive value, the percentage of correct positive PreMA predictions.

|  |  |  |  |
| --- | --- | --- | --- |
| *MENTHU alone* | 45.93% | 95.64% | 55.08% |
| *MENTHU@4 alone* | 41.53% | 97.36% | 64.72% |
| *inDelphi alone* | 23.45% | 99.15% | 76.19% |
| *MENTHU or inDelphi* | 50.49% | 95.28% | 55.46% |
| *MENTHU and inDelphi* | 18.89% | 99.51% | 81.69% |
| *MENTHU@4 or inDelphi* | 46.91% | 96.95% | 64.14% |
| *MENTHU@4 and inDelphi* | 18.08% | 99.56% | 82.84% |

**Figure 4.**
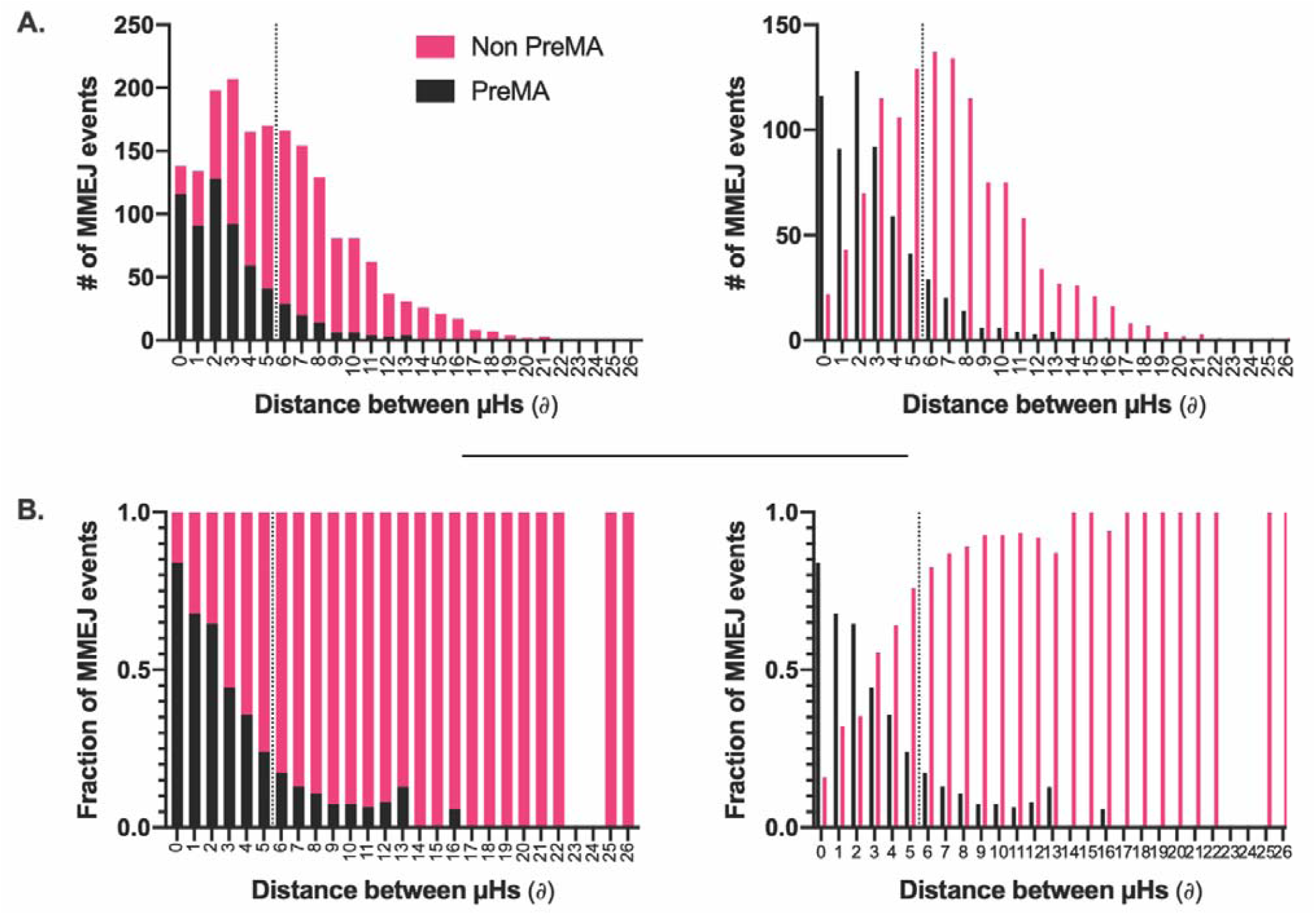
PreMA distribution of MMEJ events as a function of the distance between the microhomologies employed for repair. A. Stacked (left) and staggered (right) distributions of the number of MMEJ repair events in a large gene editing data set (19) and their PreMA status were plotted as a function of the distance between the microhomologies (µHs) used for repair (*∂*). The amount of MMEJ events increases after a *∂* of 1bp and then decreases consistently as a function of *∂* after 5bp. B. The fraction of PreMAs across each *∂* bin in A is plotted as a function of *∂*. The PreMA fraction decreases in an exponential-like fashion as a function of *∂*. The dotted lines in both A and B represent the classification threshold employed by MENTHU for PreMA predictions. Everything to the left of the dotted line is predicted as PreMA as long as the corresponding MENTHU score is ≥ 1.50.

#### Moon Rover

A logistic regression model with MENTHU score and the distance in base pairs between the µH pair for the top predicted MMEJ-based outcome as inputs and a binary PreMA/non-PreMA classification as the output.

#### Moon Walker

A gradient boosting model (25) based on decision trees (27) was trained on the same dataset as Moon Rover using the same input/output scheme. The hyperparameter combination that yielded maximum ROC area under the curve (AUC) utilized 450 trees with 6 levels of interaction depth, a minimum of 10 observations per node and a 0.01 shrinkage. ROC curves for the PreMA prediction performance on the test set by MENTHU, inDelphi, Moon Rover, and Moon Walker were generated (Figure 5). Moon Rover and Moon Walker each showed small but distinct improvements over MENTHU.

**Figure 5.**
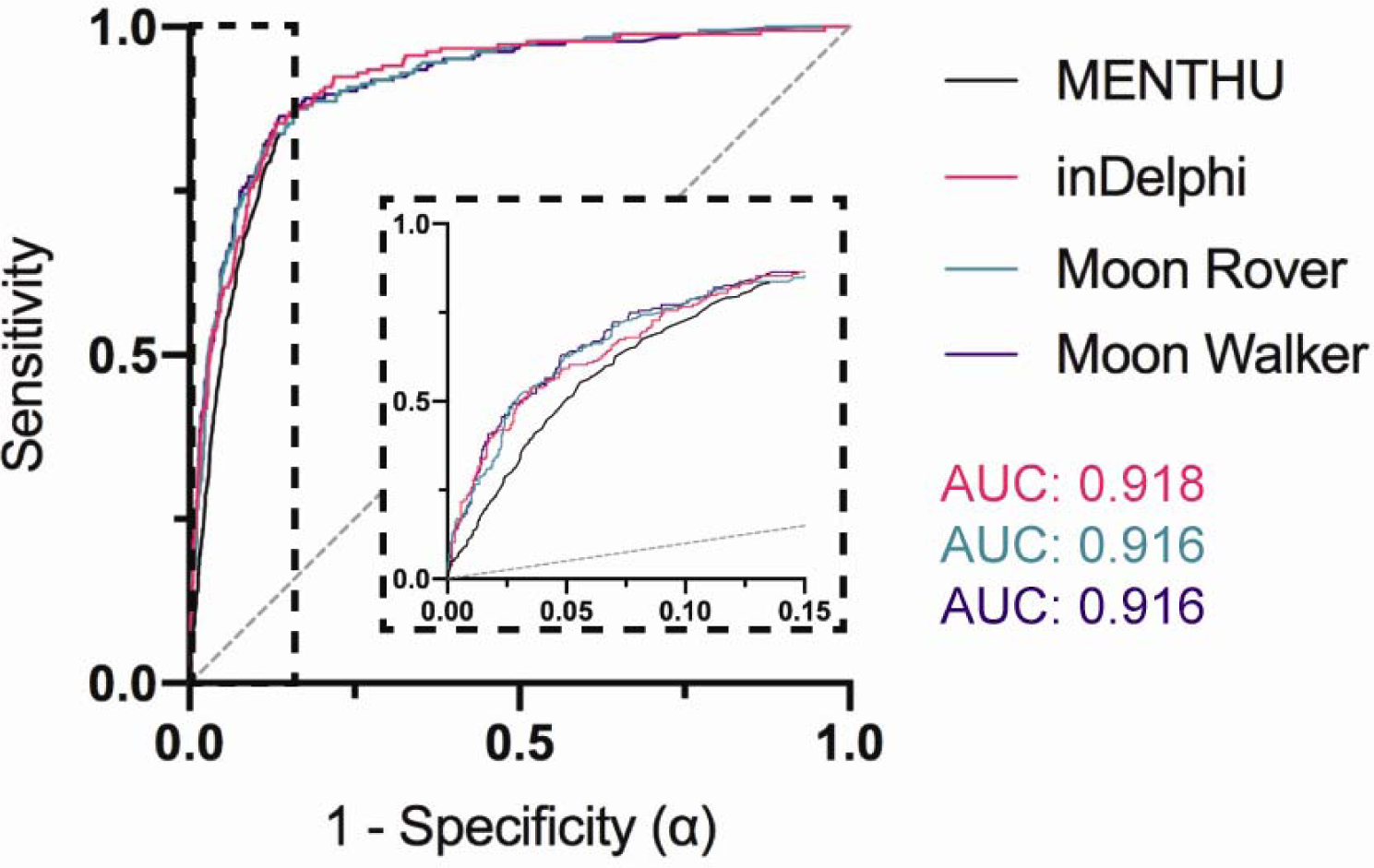
Receiving Operating Curves (ROCs) comparing the prediction performance of MENTHU and inDelphi to that of the novel MENTHU-based tools Moon Rover and Moon Walker. Moon Walker and Moon Rover are two machine-learning-based tools that utilize the same two features for PreMA predictions that MENTHU uses: the MENTHU Score and the distance between the microhomologies used for most expected MMEJ repair outcome. The ROC curves displayed represent the PreMA prediction performance of MENTHU, inDelphi, Moon Rover and Moon Walker on the out-of-sample validation set described on Figure 2D. Here, sensitivity is plotted against 1 – specificity (or the probability of a type I error: α) as a function of varying prediction thresholds. See Figure 3 legend for explanation on MENTHU and inDelphi thresholds. The inset is a blowup of the region where MENTHU is present. The area under the curve for inDelphi, Moon Rover, and Moon Walker are 0.918, 0.916, and 0.916, respectively.

### Estimation of PreMA frequency and distribution in vertebrate genomes

To assess the usefulness of PreMA targeting for reverse genetics applications, we investigated the quantity and localization of MENTHU-predicted PreMAs across 54 (28 human, 26 zebrafish) genes. The estimated PreMA frequency was consistent with previous reports (15,23), amounting to ∼10% of all targetable sites for both human and zebrafish. As expected, Cas12a, a nuclease with more targeting constraints than Cas9 (TTTV vs NGG) (28), displayed fewer potential knockout-inducing PreMAs. We also found that, when considering Cas9 alone (i.e. NGG PAMs), 81% of the genes screened (44 out of 54) has at least one predicted PreMA site in the first 30% of their coding sequences. This number increases to 88% (48/54) when also considering Cas12a.

## DISCUSSION

The success of gene editing applications from gene discovery to gene therapy is critically dependent on the specific sequence changes made at each genetic locus. For example, different outcomes due to as little as a single base difference such as in-frame versus frameshift alleles has the potential to substantially alter the observed phenotype in gene therapy uses. Similarly, a failure to generate a true loss of function allele could yield a false negative result for gene discovery testing. The goal of this study was to deploy a new tool, MMEJ-based gene editing, for improvements in gene editing precision applications (Figure 6).

**Figure 6.**
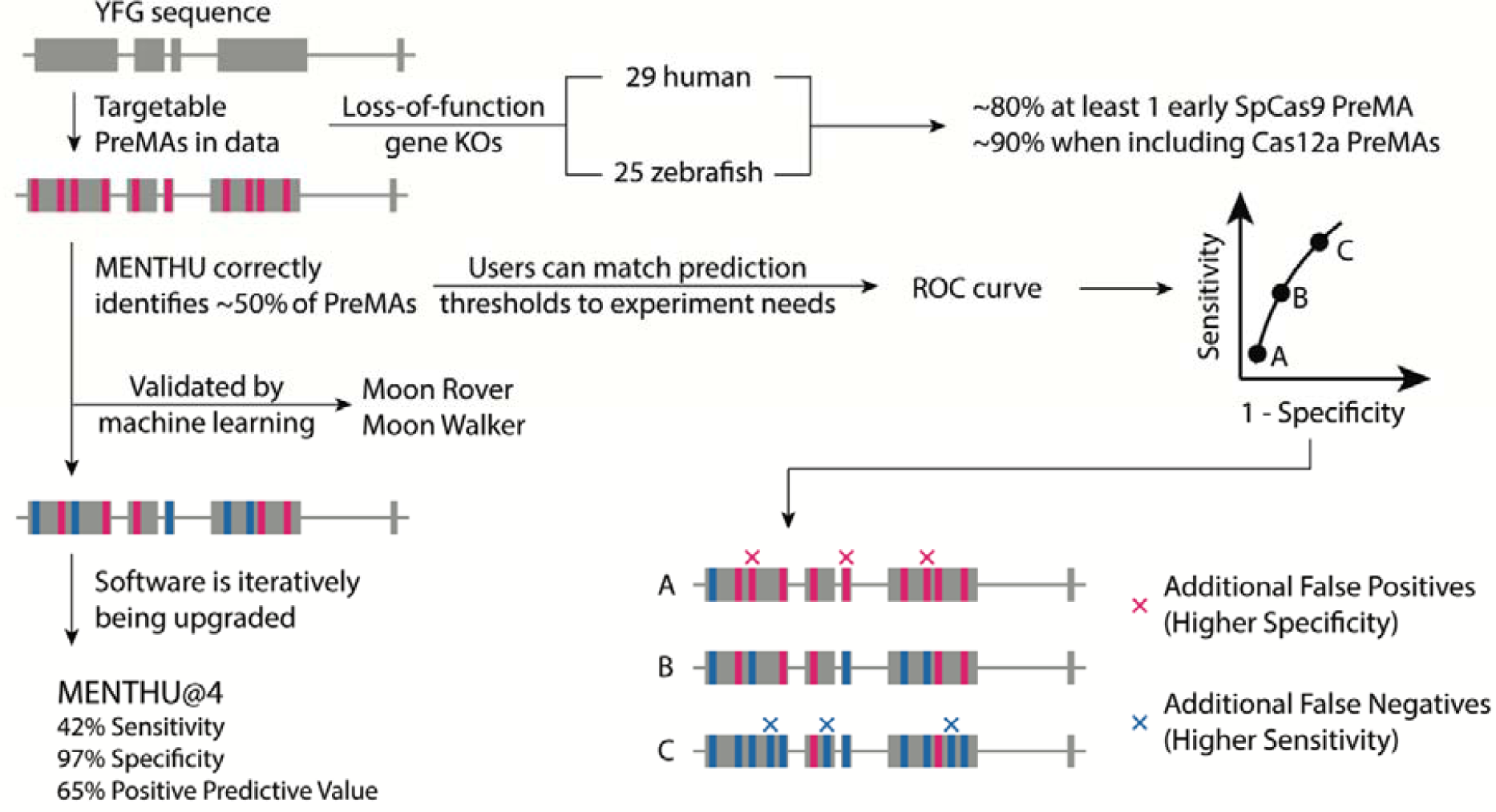
MMEJ-targeting of double-strand break sites for functional genomics and gene therapies. We sampled 54 vertebrate genes for knockout-generating PreMAs using MENTHU and estimated that the majority of vertebrate genes should possess at least one early out-of-frame PreMA. MENTHU was able to identify close to 50% of PreMAs in a large mouse embryonic stem cell line. This predictive performance was validated by comparing it to that of inDelphi, a popular tool in the field and to machine learning based models using the same features MENTHU uses for prediction. MENTHU allows users to match the prediction stringency based to the individual needs of the experiment by changing the corresponding prediction thresholds. Finally, we upgraded MENTHU to MENTHU@4 by updating its prediction parameters using a large dataset with ∼100 fold more training loci than the original version.

We have highlighted how biasing DNA repair mechanisms towards MMEJ reduces the heterogeneity of gene editing outcomes resulting from the more common NHEJ repair pathway, and thus has important advantages for reverse genetics and therapeutic applications. Here we use MENTHU and inDelphi as computational tools to identify DSBs likely to preferentially undergo MMEJ repair (PreMAs; Figure 1B), and thus potentially helpful for deploying preferentially MMEJ gene edited loci. To assess the generalizability of MENTHU results beyond the pioneering in vivo work using the zebrafish model (15), we measured the ability of MENTHU to predict PreMAs in different cellular states with an emphasis on a differential vertebrate cell type with multipotency using the large out-of-sample mESC cell gene editing dataset (20). In parallel, we also compared the mechanistically informed MENTHU algorithm against inDelphi (23), a machine learning-based tool trained to predict DSB repair outcomes in this same mESC cell type. Figure 3A explores how both tools, as initially reported, provide distinct advantages in their ability to identify PreMAs. MENTHU was able to identify close to twice as many PreMAs as inDelphi, amounting to around half of all available PreMAs in the dataset, though at a higher false positive rate. Even so, most of these false positives came from a slight overestimation of outcome frequencies (i.e. the actual repair outcome not reaching the required 50% minimum frequency for PreMA classification), and still displayed the predicted outcome sequence. This was also the case for the predicted true positives. Assuming this trend generalizes over multiple cell types, these results suggest that MENTHU could be a valuable tool for genome-wide gene discovery applications. The less sensitive inDelphi, on the other hand, does provide users with a ∼20% increase in PPV. This suggests that inDelphi could be useful as a PreMA confirmation tool for highly desirable DSBs.

However, exploring the ROC curves for both tools (Figure 3B) gives a more complete picture. First, the inDelphi results appear marginally better overall (since curves closer to the upper left in an ROC plot are better), but note that this comparison is potentially biased in favor of inDelphi since the test set comprised data from the same type of mammalian mESC cells as the inDelphi raining data (while MENTHU was trained on zebrafish cells). Also, the curves suggest that the performance differences noted above are mostly due to the specific prediction thresholds chosen for classification in the initial publications, since results for both models approximate each other by choosing different thresholds. Unfortunately, due to the nature of their repair outcome predictions, choosing a different prediction threshold in the case of inDelphi is counterintuitive. The reason is that inDelphi predicts the frequencies of the different genetic outcomes per DSB directly. As such, claiming that a threshold different to 50% should be used to observe a single repair outcome 50% or more of the time is contradictory. Additionally, inDelphi does not currently enable users to modify the prediction threshold that gives rise to their predicted frequencies. In contrast, MENTHU provides the user the ability to filter out results below a user-specified MENTHU score, enabling users to choose this threshold to their liking (22). This paper aims to guide MENTHU users so they can fully take advantage of the ability to modulate sensitivity and specificity of their predictions.

Figure 4 suggests the existence of a 0-4bp *∂* window to maximize PreMA repair outcomes. While the proportion of PreMAs decreases consistently within this window (Figure 4B), we observed a higher number of MMEJ events when *∂* ≥ 2bp. Shifting the MENTHU *∂* requirement down to only include this 0-4bp window (MENTHU@4) increased the prediction PPV and specificity without a large sacrifice in sensitivity. The ROC curves displayed in Figure 5 display the classification performance of the original MENTHU (MENTHU@5), inDelphi, and the MENTHU-based Moon Rover and Moon Walker on the same out-of-sample test set. For all levels of specificity, Moon Rover and Moon Walker achieved a higher sensitivity than MENTHU and displayed performances virtually indistinguishable from each other (AUC, area under the curve, of 0.916 for both) and from that of inDelphi (AUC of 0.918). Moon Rover and Moon Walker, therefore, provide alternatives to MENTHU with performance levels comparable to inDelphi, without any of the conceptual issues that arise when customizing the prediction threshold.

Our results suggest that the addition of PreMA-targeting schemes to experimental pipelines using MENTHU or MENTHU-like tools is beneficial for both gene therapy and genome-wide reverse genetics applications. More specifically, we have shown that MENTHU was able to identify close to half of all available PreMAs in a large dataset. By looking at the MENTHU-predicted PreMA distribution of individual genes we also aimed to investigate if PreMA targeting would be useful at an intragenic scale, particularly for gene knockout experiments. Based on the PreMA screening of 54 vertebrate genes, we estimate that, on average, four out of every five genes display at least one SpCas9-targetable PreMA within the first 30% of their coding sequences, and this becomes nine in ten genes when considering Cas12a TTTV-PAMs as well. Of note, these are likely underestimations, since we did not account for any DSBs derived from splice site targeting. Taken together, we believe that the systematic targeting of out-of-frame PreMAs to decrease the heterogeneity of the genotypic pool that results from DSB repair should be a viable strategy for almost all genes for the generation of knockouts, potentially accelerating gene discovery and gene therapy applications, and that tools like MENTHU and inDelphi currently empower genome engineers to do so.

Another feature of interest of MENTHU is the hypothesis behind the algorithm. Firstly, MENTHU only predicts PreMAs when the µHs involved in repair are relatively close. This is due to the assumption that the kinetics of the DSB repair machinery would favor a µH pair physically proximal to each other rather than distant. The exponential decrease in PreMA fraction as a function of *∂* supports this assumption (Figure 4B). Secondly, the MENTHU score was designed as a measure of the competitiveness between the µH options the repair machinery has to bridge at a given DSB. Mathematically, the MENTHU score is the ratio between the Bae et al. (16) µH pattern scores of the top two predicted outcomes repaired by MMEJ (Figure 1B’’). These pattern scores can be interpreted as the ‘strength’ of a µH pair and have been shown to correlate with the observed occurrence frequency of the corresponding MMEJ repair outcome. In addition to the proximity between the µHs used for repair, MENTHU requires the ratio of these ‘strengths’ to be 1.50 or above and interprets this scenario as a low competition state: where one µH pair is ‘stronger’ enough than the other, resulting in a higher propensity to be picked by the MMEJ repair machinery for repair. The success of MENTHU as a PreMA predictor supports this competition hypothesis, and potentially sheds some light into the function of the underlying biochemical mechanism. This is in contrast to the deep-neural-network based inDelphi which, albeit displaying comparable prediction levels to MENTHU, is difficult to interpret in terms of features or biological mechanisms, since the multiple hidden layers rapidly transform and integrate input features. That said, inDelphi predictions are a result of an ensemble of three machine learning models: two deep networks for deletions and a k-nearest neighbor scheme for insertions, and the authors include the result of the first two in the calculation of the latter, hinting at competition between deletions and insertions.

Inspired by the competition hypothesis, Moon Rover and Moon Walker employ the same two features that MENTHU uses for PreMA predictions. In Moon Rover, only the proximity criterion displayed a significant p value (< 2 × 10^−6^)), meaning that it is unlikely to have observed the results of the logistic regression by chance alone, while the MENTHU score did not show a significant p value (p = 0.468). In Moon Walker, the relative influence of each variable was calculated using the caret package in R by averaging the accuracy improvement made by each individual predictor variable at each decision split across all decision trees. According to Moon Walker, the proximity criterion was approximately 4 times more influential for accurate PreMA classification (∼80% to ∼20%). Thus, both Moon Rover and Moon Walker suggest that µH proximity is an important factor for PreMA prediction, and probably relevant to the underlying MMEJ repair biochemistry. Perhaps not as influential as proximity, the MENTHU score still improves predictions across all MENTHU-based classifiers, and future studies should investigate how to better quantify and measure µH competitiveness. The pattern scores that comprise the MENTHU score are metrics that aggregate µH length, GC content, and expected deletion length, and we are agnostic as to whether the pattern score is the best possible surrogate for µH ‘strength’. It is also likely that the features described above are not the only decisive factors in swaying the repair machinery to MMEJ preferentially, and we expect that the biochemical context surrounding the repair process to be heavily influential to factors such as cell type, DNA methylation, and more.

Precise genome engineering technology is functionally a two-step process: the generation of a specific DSB and its subsequent repair. Currently, the efficient generation of consistent DSB repair outcomes remains an important bottleneck for precision gene editing, with traditional nuclease design yielding around a 10% chance for a PreMA reagent (15,23) (for reference, 10.4% of Allen et al. (20) gRNAs are PreMA reagents). MENTHU and inDelphi represent two computational tools that offer genome engineers better control over the second step, enabling researchers to generate more consistent genotypes for their gene editing experiments. Here, we show that tools such as MENTHU and in Delphi can identify large fractions (46% and 23%, respectively) of all available PreMA sites on an independent dataset with over 50% precision (PPV). We also show that SpCas9-tagetable PreMAs likely to result in genetic KOs are found at considerable frequencies in the genome. Based on a screen of 54 vertebrate genes, we estimate the existence of at least one PreMA reagent expected to cause a truncation mutation in over 80% of the genes, with this number being closer to 90% when also considering Cas12a-targetable sites. PreMA screening thus represent a novel precision gene editing approach to facilitate consistent and reproducible outcomes for gene therapy and gene discovery applications.

## AVAILABILITY

The DNA double-strand break repair data by Allen and collaborators is available at https://figshare.com/articles/processed_mutational_profiles/7312067.

MENTHU is hosted at genesculpt.org/menthu.

inDelphi is hosted at indelphi.giffordlab.mit.edu.

## ACCESSION NUMBERS

ENSEMBL accession numbers for the CDS sequences screened for early loss-of-function PreMAs are included in Table 1.

## ACKNOWLEDGEMENT

We wish to thank the Parts Group at the Wellcome Sanger Institute for providing access to their DNA double-strand break repair dataset, Dr. Carla Mann for her work and guidance in developing MENTHU, Dr. Hirotaka Ata for the original MMEJ competition concept that inspired this project, and the members of the Ekker lab for their encouragement and support.

